# Accounting for Apparent Deviations between Calorimetric and van’t Hoff Enthalpies

**DOI:** 10.1101/210351

**Authors:** Samuel A. Kantonen, Niel M. Henriksen, Michael K. Gilson

## Abstract

**Background:** In theory, binding enthalpies directly obtained from calorimetry (such as ITC) and the temperature dependence of the binding free energy (van’t Hoff method) should agree. However, previous studies have often found them to be discrepant.

**Methods:** Experimental binding enthalpies (both calorimetric and van’t Hoff) are obtained for two host-guest pairs using ITC, and the discrepancy between the two enthalpies is examined. Modeling of artificial ITC data is also used to examine how different sources of error propagate to both types of binding enthalpies.

**Results:** For the host-guest pairs examined here, good agreement, to within about 0.4 kcal/mol, is obtained between the two enthalpies. Additionally, using artificial data, we find that different sources of error propagate to either enthalpy uniquely, with concentration error and heat error propagating primarily to calorimetric and van’t Hoff enthalpies, respectively.

**Conclusions:** With modern calorimeters, good agreement between van’t Hoff and calorimetric enthalpies should be achievable, barring issues due to non-ideality or unanticipated measurement pathologies. Indeed, disagreement between the two can serve as a flag for error-prone datasets. A review of the underlying theory supports the expectation that these two quantities should be in agreement.

**General Significance:** We address and arguably resolve long-standing questions regarding the relationship between calorimetric and van’t Hoff enthalpies. In addition, we show that comparison of these two quantities can be used as an internal consistency check of a calorimetry study.

**Highlights:**

- Agreement within ~0.4 kcal/mol between calorimetric and van’t Hoff enthalpies can be achieved for systems with typical heat and concentration errors, if solution non-ideality is not an issue.
- Concentration error chiefly affects calorimetric enthalpies, while error in measured heat chiefly affects van’t Hoff enthalpies.
- Large discrepancies between calorimetric and van’t Hoff enthalpies can be used to flag experimental error.
- There is no theoretical basis to expect discrepancies between these two methods of determining the binding enthalpy.

## 1. Introduction

The thermodynamics of molecular recognition is of central importance in a range of biological and biomedical fields, spanning molecular biophysics to drug design. The thermodynamic parameter of primary interest is the standard binding free energy, ΔG°, as it determines the binding affinity; but the standard binding enthalpy, ΔH°, and entropy, ΔS°, are also of interest, as they further characterize the molecular processes associated with binding, and can be used as reference data to test and improve the accuracy of computational molecular simulations^1^. Until about 25 years ago, the chief approach to measuring the binding enthalpy was to measure the binding free energy at several temperatures and analyze the resulting data with the van’t Hoff relation^2^,

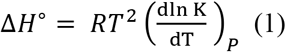

where K is the binding constant, R is the gas constant, T is absolute temperature, and the partial derivative is taken at constant pressure. The situation changed with the introduction of isothermal titration calorimetry (ITC) instrumentation^3^, which directly determines not only ΔG° but also ΔH° from measurements at a single temperature.

Somewhat unexpectedly, however, ΔH° values obtained from the van’t Hoff method and from the direct calorimetric method for the same binding reaction were early reported to be inconsistent in several cases^4–6^, with deviations of up to 10 kcal/mol. Building on these discrepancies, it was argued that the theory underlying the van’t Hoff expression might not apply in all cases, so that inconsistencies between the direct and van’t Hoff enthalpies should be expected^7,8^. However, this view was vigorously disputed^9,10^, and at least one experimental study showed that apparently large inconsistencies (up to ~3 kcal/mol) actually could be statistically insignificant^11^. Indeed, one limitation of this experimental study was the use of a relatively early model calorimeter, which lacked the precision of more modern instruments (J.R. Horn, personal communication), and thus could not provide a strong test of the consistency between the values of enthalpy from the van’t Hoff and direct calorimetric methods. More recent ITC examinations of this issue have used higher-precision instruments. In one, statistically significant deviations between van’t Hoff and direct calorimetric enthalpy values were tentatively ascribed to procedural issues, rather than to inapplicability of the van’t Hoff relation^6^. In another, though, a new apparent experimental violation of the van’t Hoff relation was demonstrated experimentally, and it was again argued on theoretical grounds that consistency between the two values of the binding enthalpy should not be expected^12^. However, both of these more recent studies involved injection of dications, Ba^2+^ or Ca^2+^, into the reaction cell, an experimental design that could conceivably have generated temperature-dependent changes in activity coefficients and partial molar enthalpies that might complicate interpretation of the data.

Here, we reexamine this issue with new experimental tests of the consistency between van’t Hoff and directly measured enthalpy values, for ITC experiments that reduce the potential for changes in solution ideality during the experiment by using an electrically neutral molecular host, β-cyclodextrin, as the titrant. Such compact molecular recognition systems are finding increasing application as models to evaluate and improve the reliability of computational models of binding^13–15^, and we are particularly interested in assessing the reliability of binding enthalpies, because computational methods of obtaining these from simulations now offer the possibility of comparing not only binding free energies, but also enthalpies, with experiment^14,15^. In addition, a number of prior studies that have reported discrepancies between van’t Hoff and direct calorimetric enthalpies also focused on host-guest systems, such as crown ethers and cyclodextrins^4–6,12^. However, the basic results are expected to be applicable not only to other host-guest systems, but also to biomolecules. The experiments are carried out on a modern instrument with a low level of uncertainty in its heat measurements^16^. We furthermore use mathematical modeling to investigate how two common sources of experimental error, concentration error and heat error^16^, can generate apparent violations of the van’t Hoff relation. We find that, although van’t Hoff enthalpies are sensitive to errors in the binding free energy, they may nonetheless be expected to agree with directly measured enthalpies to within about 10%, given typical experimental errors. We also find that concentration error chiefly affects the direct enthalpy, while heat error chiefly affects van’t Hoff enthalpies, and that marked inconsistency between the van’t Hoff and direct enthalpies can be used to flag pathological measurements. We close by discussing the implications of these results and considering recent theoretical arguments against the applicability of the van’t Hoff relation to aqueous binding thermodynamics.

## 2. Materials and Methods

### 2.1 Materials

β-cyclodextrin (catalog no. C-4767), rimantadine hydrochloride (catalog no. 390593), and amantadine hydrochloride (catalog no. 138576) were obtained from Sigma-Aldrich Company (St. Louis, MO). Previous older batches (over one year after purchase) of β-cyclodextrin showed some evidence of aggregation in newly made solutions that had stood for on the order of an hour or more, based on slight clouding of concentration solutions and spectrophotometric detection, so a new lot was used for experiments presented in this study. Solutions made from the new lot showed no visible evidence of aggregation. NMR spectra were taken for β-cyclodextrin, rimantadine, and amantadine to verify structure and purity.

### 2.2 Isothermal Titration Calorimetry Experiments

ITC experiments were performed using a MicroCal model VP-ITC (MicroCal, Northampton, CT, Serial Number 01-08-930). Previous work has emphasized that even minor deflections in the power baseline during ITC experimentation can lead to nontrivial errors that, when feasible, requires detailed analysis to correct. We therefore investigated sources of baseline noise for our experimental setup. For our VP-ITC instrument, we found that small, deliberate vibrations of the laboratory bench on which the calorimeter was set caused significant deflections in baseline. To prevent this, the calorimeter was placed in an isolated room on a 2” thick block of urethane foam. This setup eliminated deflections during deliberate vibration of the bench and provided a more stable baseline. Additionally, a purpose-built, clear acrylic shield was used to reduce possible temperature shifts due to drafts. The instrument’s built-in Y-axis calibrations were performed, at 27 C**°**, to verify that the instrument was responding to known power inputs within the 1% error tolerance prescribed by the manufacturer.

Solutions of β-cyclodextrin, rimantadine, and amantadine were prepared in 10 mM phosphate buffered saline (pH 7.4). In all experiments, β-cyclodextrin was titrated into the cell, in order to take advantage of both its negligible heat of dilution (Supplementary Figure 1) and its electrical neutrality, which reduces the likelihood of large changes in activity coefficients and partial molar enthalpies in the course of the experiment. Because the measured binding enthalpy is particularly sensitive to the concentration of the syringe titrant, the β-cyclodextrin solutions were prepared in relatively large quantities, as this reduces errors in the concentration due to weighing errors. The solutions thus were prepared in 250 mL volumetric flasks at concentrations of 12.8-13.4 mM, which corresponds to about 3 grams of β-cyclodextrin per 250 mL batch of solution. The concentration of both guests was 1.1-1.4 mM. This is the maximum concentration of β-cyclodextrin attainable in aqueous buffer. Solutions of the guest molecules in the ITC cell were prepared in 25 mL volumetric flasks. Masses of both host and guest were measured using a Sartorius CPA225D Micro Balance. An additional set of experiments for each guest was performed with reduced concentrations (0.27 mM rimantadine and 0.23 mM amantadine, with 5.1-6 mM β-cyclodextrin), using the same procedures described above, to check for evidence of nonideal solution properties. For each guest, duplicate experiments were performed, and fresh solutions were prepared for each replicate. However, a single set of cell and syringe solutions was used for each complete temperature series, to keep the concentrations precisely the same across the measurements in the series.

ITC experiments were designed to allow maximum signal while staying within the instrument’s limitations in measured power, and thus to maximize the signal-to-noise ratio. Thus, at each temperature, the reference power was set as low as possible without allowing overshoot due to generation of binding heat at a rate in excess of the reference power. For amantadine, 27 10-uL injections were performed at each temperature. For rimantadine, runs were done using 54 5-uL injections and 27 10-uL injections. Smaller injection sizes were chosen for rimantadine because of its more enthalpic reactions; for the concentrations chosen for rimantadine, the raw heat signal from *10 µ*L injections overwhelmed the calorimeter, causing problems with the baseline. For all experiments, the first injection was discarded; this practice is ubiquitous in ITC experimentation, as the heat of the first injection is often inappropriately small, due to diffusive mixing of cell and syringe solutions within the tip of the syringe prior to the first injection. The time between injections for a typical experiment in this study was extended from the default of 120 seconds to 300 or 500 seconds, to allow for a more complete return to baseline.

### 2.3 Generation of Artificial ITC Data

We generated and analyzed artificial Wiseman plots based on thermodynamic parameters for the reversible association of a model system previously studied by both our laboratory and others: the adamantyl-based drug rimantadine with the cyclic oligosaccharide β-cyclodextrin. The data were modeled for our own experimental setups with rimantadine in the cell and cyclodextrin in the syringe, with binding thermodynamics and Wiseman plot C values similar to experimental conditions from 300 K to 340 K (50 to 6, respectively). All modeled data use 25 injections of 10 *µ*L to generate Wiseman plots. Following our prior investigation into heat error for this instrument^16^, we estimated the standard deviation of the measured heat of a given injection as

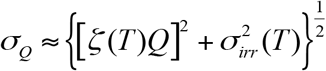

where *σ*_*Q*_ is the standard deviation of the measured injection heat; ξ(*T*) is the temperature dependent coefficient of the proportional error component; and *σ*_*irr*_(*T*) is the standard deviation of the temperature-dependent irreducible heat noise. Based on our prior study^16^, we set ξ to 0.01 (1%) for T=27-57C, and to 0.03 (3%) for experiments at 67C. Similarly, σ_irr_ was set to 0.13 *µ*cal and 0.39 *µ*cal for experiments at 27-57C and at 67C, respectively.

We also used model ITC data to determine whether thermal expansion of the syringe and cell solutions, as T is increased from 300K to 340K to generate data for the van’t Hoff method, might lower concentrations enough to materially affect ΔH_Direct and_ ΔH_VH_. Accounting for the ~1.4% thermal expansion over this temperature had very little effect on the results (Supplementary Table 2), so we did not include thermal expansion in subsequent modeling or data analysis.

### 2.4 Analysis of ITC Data

The Origin 7.0 software was used to process the raw experimental data by integrating it to give the heat release per injection. Using custom Python scripts, either artificial model data or the experimental raw heats were normalized by concentration of injectant to produce a Wiseman plot, and values of K and ΔH for the binding reactions were determined by using non-linear optimization (Marquardt-Levenberg) with theoretical equations described by Wiseman^3^. The values of the binding enthalpy from this procedure are termed the direct results, ΔH_Direct_, while values of the binding enthalpy obtained from the van’t Hoff relationship (see below) are termed van’t Hoff results, ΔH_VH_. (Although the superscript “o” is omitted, for simplicity, both quantities pertain to the usual standard condition where the reactants and product are present in a hypothetical ideal 1 M solution.)

We used bootstrap analysis to study how heat error and concentration error propagate to the fitted quantities ΔH_Direct_ and ΔH_VH_, for both the experimental and simulated data. For each point on the Wiseman plot, heat error was modeled by sampling the heat of each injection from a Gaussian with a mean of the measured or simulated heat, and standard deviation σ_Q_, from the equation above. These Gaussian samples allowed the construction of multiple artificial Wiseman plots, each with its data points sampled from its respective Gaussian. Additionally, each artificial Wiseman plot was recalculated multiple times with various concentration errors assumed. Importantly, the same concentration was always used for each full set of Wiseman plots across temperatures, to simulate doing a set of temperature points measured for the same solution, as done in our experiments. Thus, although there is some error in the concentration of the solution, the same solution is used at all temperatures to get an internally consistent set of binding free energies for use in the van’t Hoff analysis.

Values of K and ΔH_Direct_ were fitted to each resulting Wiseman plot, to yield a statistical distribution of these fitted quantities, allowing calculation of the mean and standard deviation of K and ΔH_Direct_ across the bootstrapped samples. This bootstrapping can more fully capture the uncertainties in the reported thermodynamic quantities than the common approach, used in Origin and other programs, of reporting errors based only on the fit to a single Wiseman plot^16^. The overall mean and SEM of the ΔH_VH_ and K values for each temperature point, across experimental duplicates, were determined by taking the mean and standard deviation over all replicates, from bootstrapping and from the duplicated measurements. Thus,

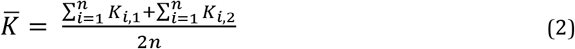

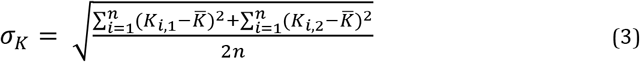

where *K*_*i,1*_ is the fitted value from the *i* ^th^ Wiseman plot during bootstrapping, over n Wiseman plots based on the first measurement, and *K*_*i,2*_ is the fitted value from the *i* ^th^ Wiseman plot during bootstrapping, over n Wiseman plots based on the duplicate measurement. The mean and standard deviation of ΔH_VH_ was determined with analogous formulae. This bootstrapping process realistically models the propagation of uncertainties in the raw data to the derived thermodynamic properties. For model Wiseman plots, the data were processed both with the stoichiometry, N, fixed at 1, since one-to-one binding was used to generate the simulated data; and with N allowed to float as an additional adjustable parameter in the nonlinear optimization.

We also obtained fitted values for the change in heat capacity on binding, *ΔC*_*P*_, either by taking the slope of the direct binding enthalpies (direct method), or by taking the average of the fitted *ΔC*_*P*_ over five temperatures from the van’t Hoff method (described below). This allowed an additional comparison of direct and van’t Hoff method calculations.

### 2.5 Van’t Hoff method

The van’t Hoff enthalpies, ΔH_VH_ were computed by applying the van’t Hoff relationship to a series of binding free energies, ΔG(T), at several different temperatures, T. We used an integrated form of the van’t Hoff relationship, which is sometimes termed the Gibbs-Helmholtz equation^17^:

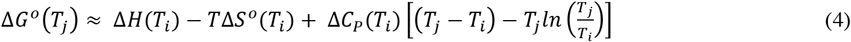

Here, *T*_*t*_ and *T*_*j*_ are two temperatures, Δ*G*° and Δ*S°* are the standard binding free energy and entropy at standard concentration (1M), Δ*C*_*p*_ is the change in heat capacity on binding, and the thermodynamic quantities are written as explicit functions of temperature. Note that the approximation in Eq 4 becomes an equality when the heat capacity is independent of temperature. For each host-guest pair, ITC experiments were run at five temperatures, *T*_*i*_, *i* = 1,2,3,4,5, and the binding free energy at each temperature was computed as *ΔG°*(*T*_*j*_) = −*RTln K*(*T*_*j*_), where *K*(*T*_*i*_) is the measured binding constant. The van’t Hoff enthalpy at temperature *T*_*i*_,Δ*H*_*vh*_(*T*_*i*_), was then assigned by using nonlinear optimization to adjust the three quantities *ΔH*_*vh*_(*T*_*i*_), Δ*S°*(*T*_*i*_) and Δ*C*_*p*_(*T*_*i*_) so that the values of *ΔG°*(*T*_*j*_) computed by Eq 4 across all five temperatures were best fit to their corresponding experimental values. Weighted, nonlinear fitting was used to account for the uncertainties in Δ*G°*(*T*_*i*_) in obtaining the van’t Hoff enthalpies.

We found that setting each value of *T*_*i*_ as a reference temperature to compute Δ*H*_*VH*_(*T*_*i*_) in this way yielded better agreement with the direct enthalpies than an alternative procedure in which only one value of *T*_*i*_ was used as a reference temperature for all five values of Δ*H*_*VH*_ (*T*_*i*_). In this manner, Δ*C*_*p*_ is fit at each temperature, with the reported Δ*C*_*p*_ being the average value across all temperatures. We also find that the most reliable van’t Hoff enthalpies are typically obtained from temperature-dependent free energies computed with the stoichiometry, N, allowed to float (see Results), so this procedure is used for all van’t Hoff enthalpies reported here, except as otherwise noted.

Experimental uncertainties in the van’t Hoff enthalpies were obtained by a bootstrapping procedure similar to that used for the direct enthalpies. For each set of bootstrapped Wiseman plots (five Wiseman plots corresponding to a given concentration error), the fitted Δ*G°*(*T*_*j*_) values from each plot were used to compute Δ*H*_*VH*_(*T*_*i*_). Thus, for each set of Wiseman plots a corresponding set of Δ*H*_*VH*_(*T*_*i*_) values is generated, and the resultant distribution of these is used to determine the mean and standard deviation.

### 2.6 Statistical measures

We wish to quantify the deviations between direct and van’t Hoff enthalpies under various conditions and with various assumptions regarding experimental error (see Results). To do this, we use procedures described above to generate *N*_*sample*_ artificial sample datasets with the desired levels of error, where each dataset includes results at *N*_*temp*_*=5* temperatures (see above), and to use nonlinear fitting to derive values of direct and van’t Hoff enthalpy from these artificial data. We then define the discrepancy, *D*, between the direct and van’t Hoff enthalpies as their mean unsigned difference:

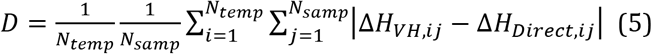

Except as otherwise noted, we allow the stoichiometry, N, to float when computing values of ΔG^o^ for use in obtaining the van’t Hoff enthalpies (see above). However, we consider allowing N to float or be held fixed at 1 when computing the direct enthalpies, and examined how this choice affected the discrepancy *D*. To quantify this effect, we define *ΔD= D*_*float*_*-D*_*fix*_ where the two values of D are computed with values of *ΔH*_*Direct*_ obtained with N allowed to float and N held fixed, respectively.

We similarly quantified the error in direct enthalpies computed by fitting to model calorimetry data with experimental error, relative to the nominal or “true” error-free values used to generate the data, as their mean unsigned error:

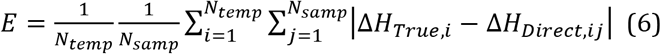

We also analogously defined *ΔE=E*_*float*_*-E*_*fix*_, which is positive if computing the direct enthalpies with N allowed to float leads to higher errors than treating N as fixed, and is negative in the opposite case.

### 2.7 Code availability

The code used to generate and analyze model data is available at (https://github.com/GilsonLabUCSD/Isothermal-Titration-Calorimetry). It allows one to generate model ITC data, based on assumed binding free energies and enthalpies over a temperature range, apply estimated concentration error and heat noise, use statistical sampling to generate model enthalpograms, and use these to obtain values of the direct and van’t Hoff enthalpies. It can thus be used to estimate the level of van’t Hoff consistency one should expect for a given binding system and estimated levels of heat and concentration error.

## 3. Results

We first report experimental ITC measurements of the binding thermodynamics of two adamantyl based guests to beta-cyclodextrin at multiple temperatures. Good agreement is obtained between the direct (ΔH_Direct_) and van’t Hoff (ΔH_VH_) binding enthalpies obtained from the same data. We then use simulated data to investigate the consequence of two key sources of experimental error, heat error and concentration error^16^, for the consistency between ΔH_Direct_ and ΔH_VH_. We examine the level of consistency expected under typical experimental conditions, and provide evidence that lack of consistency can serve as an indicator of high experimental error.

### 3.1 Experimental van’t Hoff consistency

To ascertain the level of van’t Hoff consistency that could be observed experimentally with a modern calorimeter and appropriate experimental design, we used ITC to study the binding thermodynamics of two host-guest systems, beta-cyclodextrin with amantadine and rimantadine. These experiments avoid potential technical pitfalls, because all reactants are reasonably soluble, and because beta-cyclodextrin, placed in the syringe, has a minimal heat of dilution (Supplemental Figure 1) and carries no electric charge. It is worthwhile to note that even the ionic guests give negligible heats of dilution (Supplemental Figure 1). It is reasonable to assume then, that most of the error in the measurements will stem from heat and concentration error, which we have characterized experimentally^16^.

For both host-guest pairs, measurements were taken at five temperatures, ranging from 300 K to 340 K. The average free energies and direct enthalpies of binding over two sets of replicates for both guests are presented in Table 1, with errors assigned as described in Methods. It is worth noting the clear changes in these enthalpograms as temperature increases (Figure 1): reduced sharpness of the sigmoidal Wiseman plots indicate drops in affinity, and, for rimantadine, the injection peaks are clearly taller at high temperature, indicating a more favorable binding enthalpy. This combination of changes implies entropy-enthalpy compensation, as borne out by the fitted data in Table 1. Note that these trends in free energy and enthalpy with T cannot be attributed to concentration error, because the same solutions were used across the range of temperatures in all cases. It is also worth noting that the results for rimantadine at 300K agree with previously published binding measurements for this host-guest pair to within 0.1 kcal/mol^18^.

**Table 1:** Binding enthalpies and free energies (kcal/mol) for amantadine and rimantadine with beta-cyclodextrin, from 300 to 340 K. Uncertainties are one standard deviation, generated from bootstrapping during fitting as described in Methods. For amantadine, the C values range from 2 to 10 over the temperatures series; for rimantadine, the C values range from 10 to 50.

|  | Amantadine |  |  | Rimantadine |  |  |
| --- | --- | --- | --- | --- | --- | --- |
| | $\Delta H$ | $\Delta G^\circ$ | $T\Delta S$ | $\Delta H$ | $\Delta G^\circ$ | $T\Delta S$ |
| 300 K | $-5.29 \pm 0.04$ | $-5.29 \pm 0.01$ | $0.00 \pm 0.04$ | $-6.75 \pm 0.04$ | $-6.23 \pm 0.01$ | $0.52 \pm 0.04$ |
| 310 K | $-6.08 \pm 0.06$ | $-5.27 \pm 0.01$ | $0.81 \pm 0.06$ | $-7.56 \pm 0.04$ | $-6.20 \pm 0.01$ | $1.36 \pm 0.04$ |
| 320 K | $-6.55 \pm 0.07$ | $-5.27 \pm 0.01$ | $1.28 \pm 0.07$ | $-8.34 \pm 0.05$ | $-6.19 \pm 0.01$ | $2.15 \pm 0.05$ |
| 330 K | $-7.42 \pm 0.10$ | $-5.18 \pm 0.01$ | $2.24 \pm 0.10$ | $-8.98 \pm 0.06$ | $-6.11 \pm 0.01$ | $2.87 \pm 0.06$ |
| 340 K | $-8.13 \pm 0.41$ | $-5.16 \pm 0.05$ | $2.97 \pm 0.43$ | $-9.67 \pm 0.21$ | $-6.01 \pm 0.02$ | $3.66 \pm 0.23$ |

**Figure 1:**
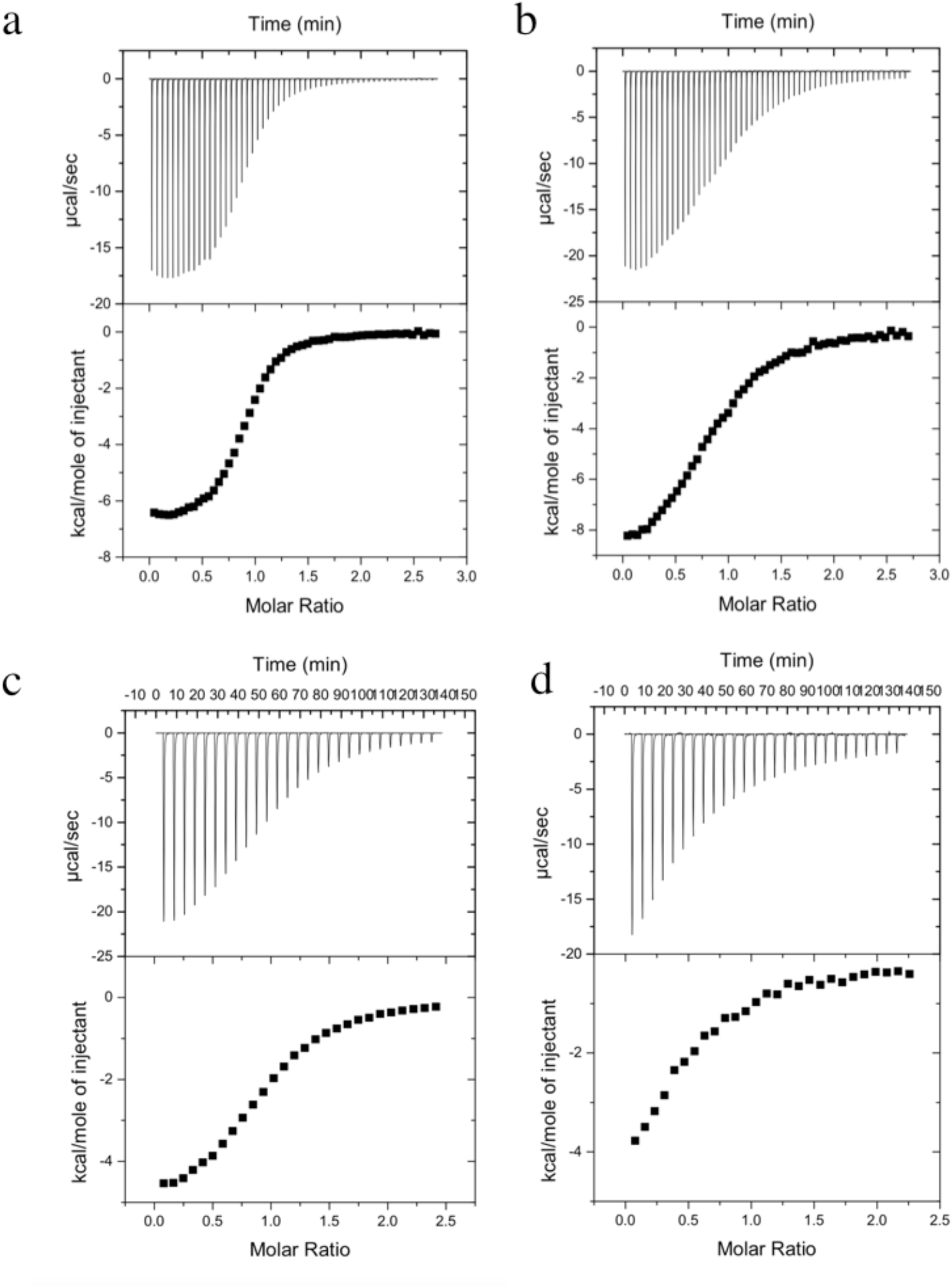
Sample experimental plots for beta-cyclodextrin binding rimantadine (a,b) and amantadine (c,d). Panels a and c show measurements at 300 K, panels b and d show measurements at 340 K. For each panel, the top half corresponds to the raw enthalpogram and the bottom to integrated enthalpograms (Wiseman plots). For each guest compound, the injectant concentrations and volumes are the same at 300K and 340K, as are the cell concentration. The ITC C values for these experiments are as follows: 50 (a), 6 (b), 10 (c), 3 (d)

The direct enthalpies are within 95% confidence intervals (about two standard deviations) of van’t Hoff enthalpies obtained from the same data (Table 2, Figure **2**), with mean unsigned deviation between van’t Hoff and calorimetric enthalpies of D=0.16 kcal/mol and D=0.39 kcal/mol for amantadine and rimantadine, respectively. However, the slopes of the van’t Hoff enthalpies with T deviate somewhat from those of the direct enthalpies. This pattern is suggestive of heat error somewhat in excess of our estimates, rather than concentration error, as indicated in Figure 4, panels b,d, and associated text (below). The heat capacity changes, ΔC_p_, derived from the two methods also agree reasonably well; the values for amantadine are 74.6 and 70.1 cal/mol/K, for direct and van’t Hoff, respectively; for rimantadine, the corresponding results are 72.7 and 93.9 cal/mol/K. The heat capacities come from the second derivative of the binding free energies with respect to temperature, and thus are more sensitive to noise than the enthalpies. Additional measurements performed with reduced concentrations of both host and guest (Supplementary Figure 3) are consistent with the present results. This indicates that the binding thermodynamics are independent of concentration in this range, and thus that the solutions used are close to ideal.

**Table 2.** Direct and van’t Hoff binding enthalpies (kcal/mol) for amantadine and rimantadine binding beta-cyclodextrin from 300 to 340 K. Errors shown are one standard deviation, generated from bootstrapping, as outlined in the Methods section.

|  | Amantadine |  | Rimantadine |  |
| --- | --- | --- | --- | --- |
| | $\Delta H_{\text{Direct}}$ | $\Delta H_{\text{VH}}$ | $\Delta H_{\text{Direct}}$ | $\Delta H_{\text{VH}}$ |
| 300 K | $-5.29 \pm 0.04$ | $-5.01 \pm 0.58$ | $-6.75 \pm 0.04$ | $-5.97 \pm 0.43$ |
| 310 K | $-6.08 \pm 0.06$ | $-5.80 \pm 0.25$ | $-7.56 \pm 0.04$ | $-6.97 \pm 0.22$ |
| 320 K | $-6.55 \pm 0.07$ | $-6.56 \pm 0.27$ | $-8.34 \pm 0.05$ | $-7.92 \pm 0.15$ |
| 330 K | $-7.42 \pm 0.10$ | $-7.29 \pm 0.59$ | $-8.98 \pm 0.06$ | $-8.84 \pm 0.33$ |
| 340 K | $-8.13 \pm 0.41$ | $-8.00 \pm 0.93$ | $-9.67 \pm 0.13$ | $-9.73 \pm 0.53$ |

**Figure 2:**
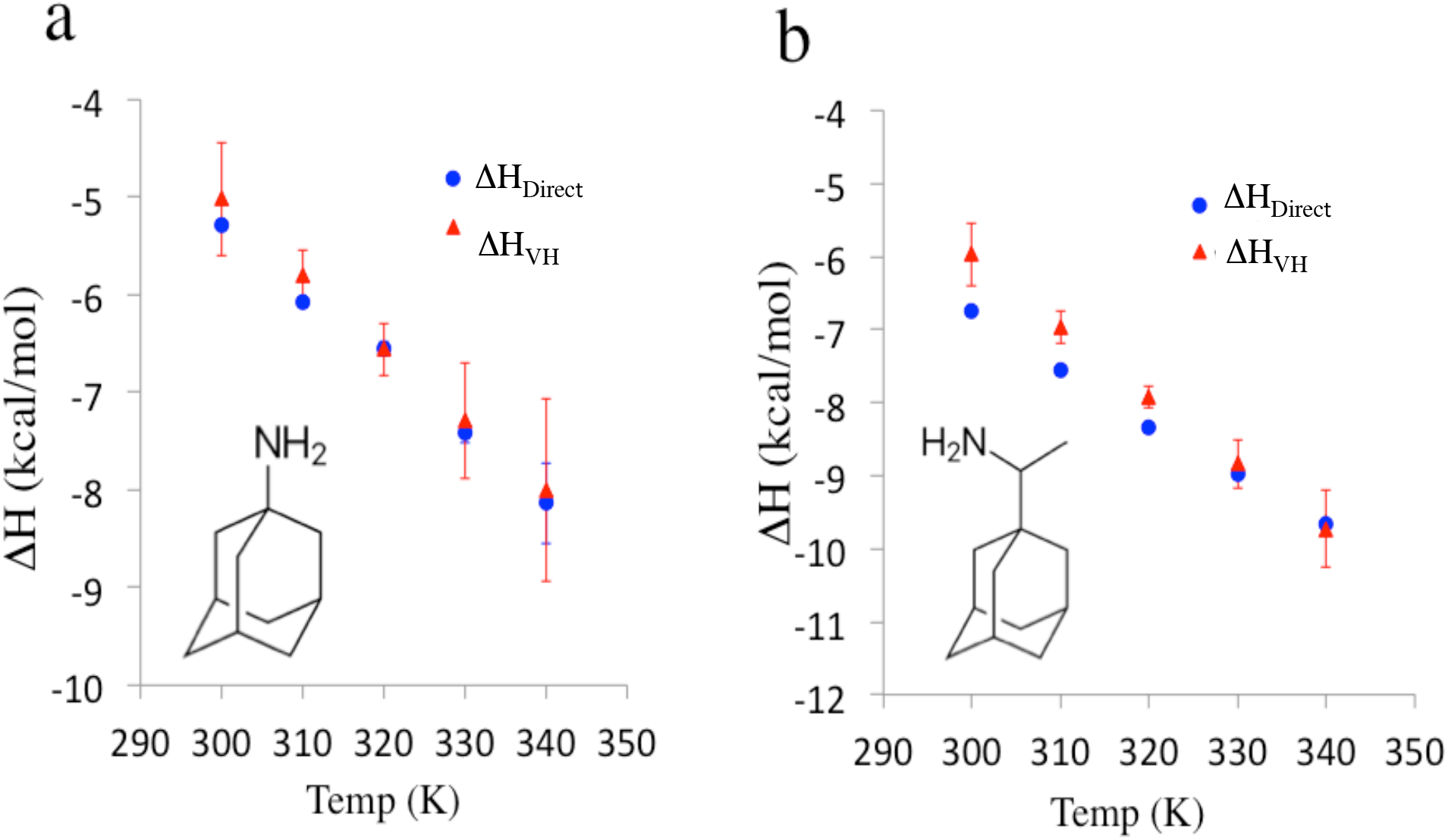
Measured direct and van’t Hoff binding enthalpies versus temperature. a) beta-cyclodextrin and amantadine. b) beta-cyclodextrin and rimantadine. Error bars shown are one standard deviation, generated from bootstrapping, as outlined in the Methods section.

### 3.2 Propagation of experimental error to calorimetric and van’t Hoff enthalpies

Here, we use simulated data to characterize the distinct ways in which concentration error and heat error propagate to ΔH_Direct_ and ΔH_VH_, to test whether severe inconsistency between these two quantities could be useful to flag data with high errors, and to estimate the level of agreement between these quantities to be expected under typical conditions for near-ideal solutions. Ideal Wiseman plots at five temperatures were generated for values of K, ΔH°, and ΔC_p_ similar to those of rimantadine binding beta-cyclodextrin (Supplementary Table 1), and assuming 1:1 binding (N=1) (Figure 3). As a methodological check, we confirmed that fitting to these ideal Wiseman plots data yielded values of ΔH_Direct_ and ΔH_VH_ that agree essentially perfectly with each other and with the binding enthalpy used to generate the model data (data not shown). We then added heat error or concentration error to each ideal Wiseman plot. In order to simulate a series of ITC measurements using the same solutions across different temperatures, the same erroneous concentrations were used across each temperature series, though each temperature series used a different assumed concentration error. Bootstrap resampling was used to generate 1000 such independent temperature series, resulting in 1000 values of ΔH_Direct_ and ΔG° for each temperature point. The values of ΔG° were furthermore used to fit 1000 van’t Hoff enthalpies at each temperature, based on Eq 4. The magnitude and character of both the concentration error and heat error used here are chosen to reflect typical experimental conditions, as determined in our prior empirical study^16^.

**Figure 3:**
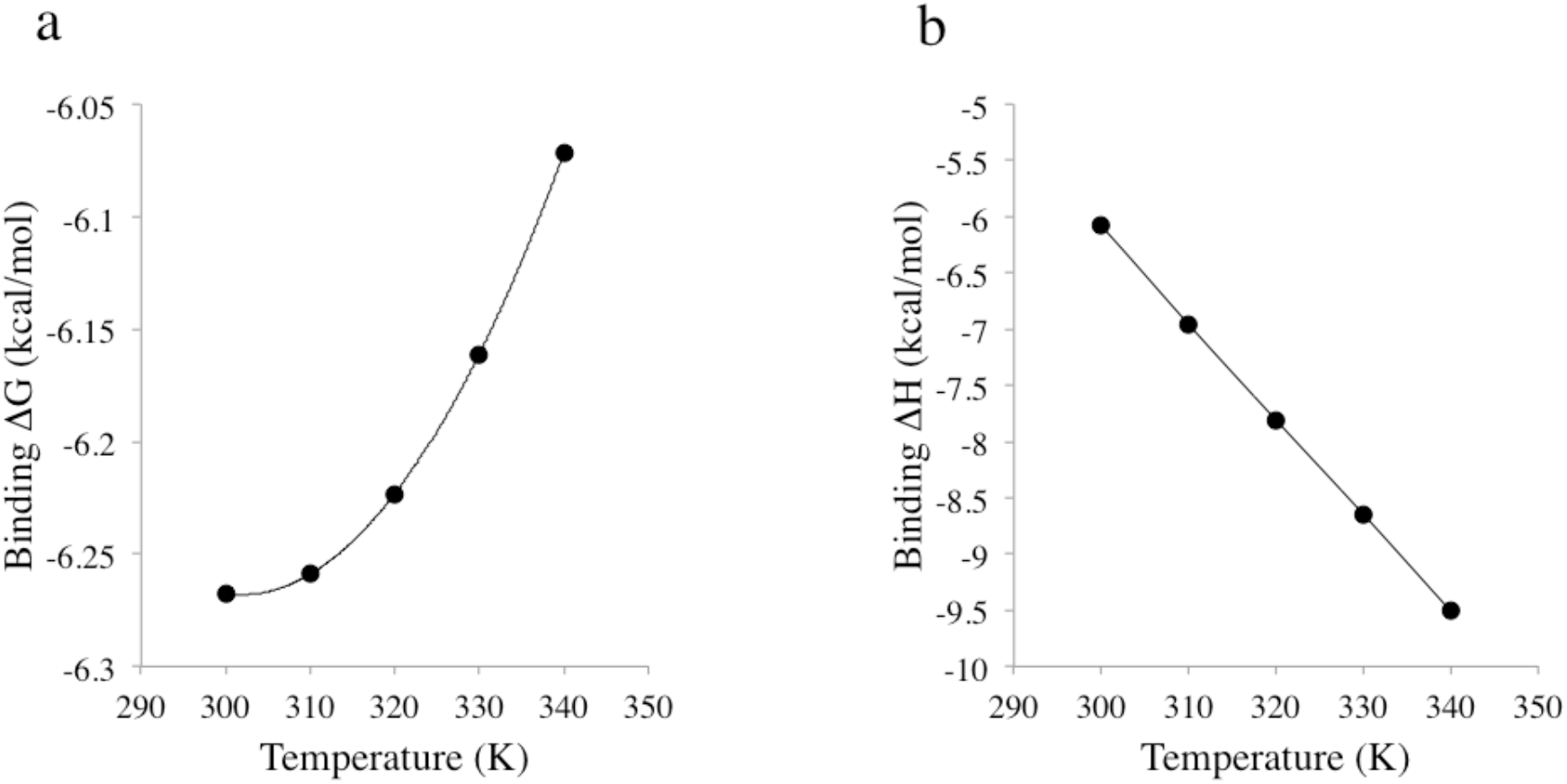
Ideal binding free energies and enthalpies, corresponding to ideal Wiseman plots. (A) Binding free energies from ideal Wiseman plots at five temperature points (B) Binding enthalpies from ideal Wiseman plots at five temperature points.

A central observation of this work is that realistic levels of concentration error and heat error lead to strikingly different patterns of error in ΔH_Direct_ and ΔH_VH_, as illustrated by representative data drawn from the 1,000 bootstrapped datasets for N floating (Figure 4) and N fixed at 1 (Figure 5). With N floating, pure concentration error propagates significantly to ΔH_Direct_ (Fig 4a), but negligibly to ΔH_VH_ (Fig 4b). That pure concentration error has minimal effect on the van’t Hoff enthalpies presumably derives from our assumption that the same solutions are used across all temperatures, as this leads to a small and nearly constant shift in ΔG° across T, which has little effect on the slope and hence on ΔH_VH_. Conversely, with N floating, heat error propagates only slightly to ΔH_Direct_ (Fig 4c), but propagates strongly to ΔH_VH_ (Fig 4d), leading to plots of van’t Hoff enthalpy versus temperature whose slopes deviate from their true value. It is worth noting that the realistic levels of concentration and heat error used here lead to errors in fitted ΔG^o^ on the order of only hundredths of a kcal/mol (Supplementary Figure 1a, b), smaller than those previously assumed in studying uncertainties in van’t Hoff binding enthalpies^19^.

**Figure 4.**
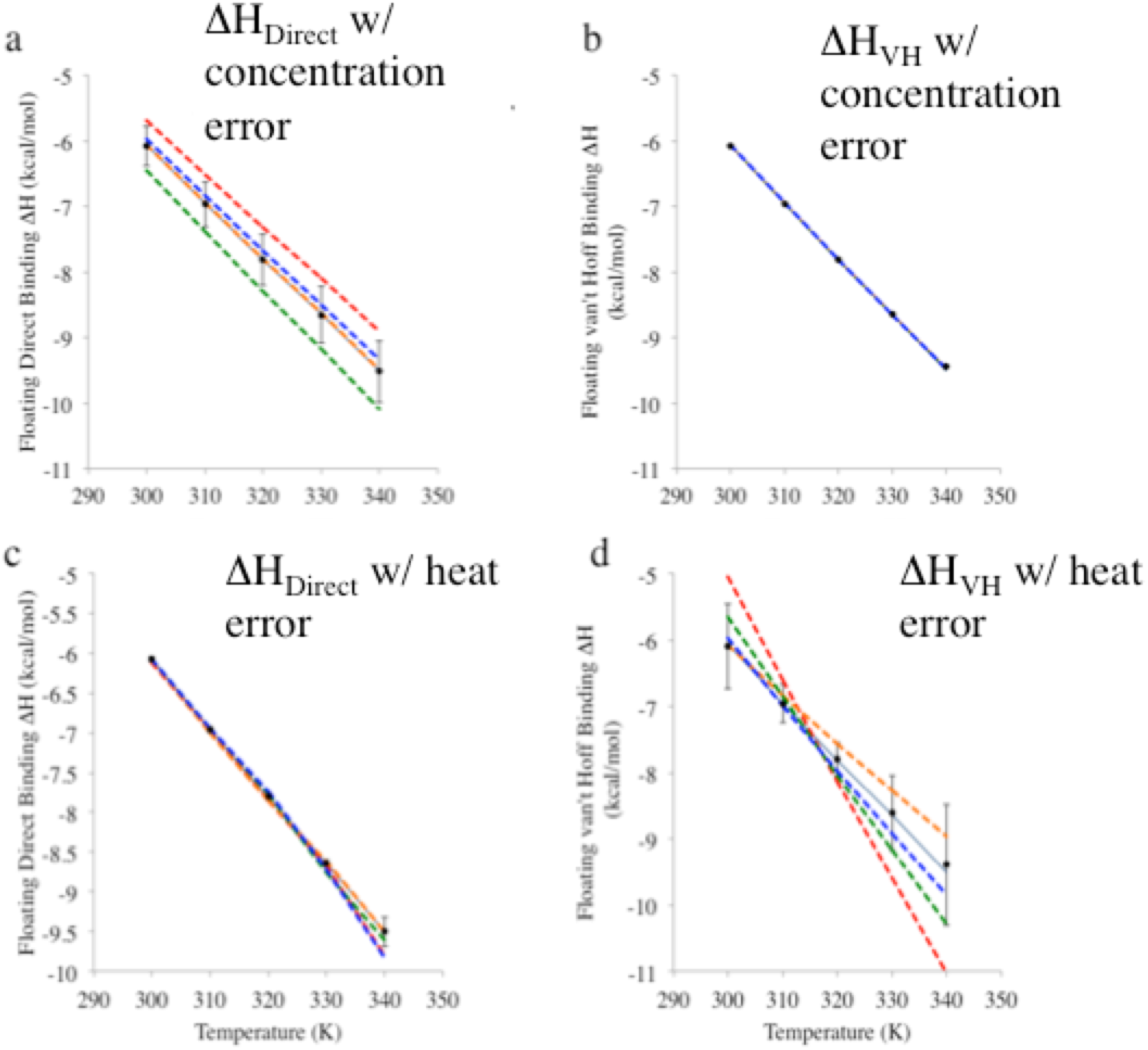
Effect of heat and concentration error on direct and van’t Hoff binding enthalpies based on binding free energies fitted with stoichiometry parameter N floating. The solid black line in each panel represents the nominal binding enthalpies used to generate the fitted Wiseman plots, while each of the dashed, colored lines plot the binding enthalpies from one set of replicates; five examples of these are included in each panel. Error bars represent one standard deviation over 1000 replicates. For concentration error, *5%* syringe and cell error was applied. For heat error, for 300-330 K points, 1% heat error and 0.13 ucal irreducible baseline error was added; for 340 K points, 3% heat error and 0.39 ucal irreducible baseline error was added. (A) Fitted direct binding enthalpy at each temperature point with concentration error applied (B) Fitting van’t Hoff binding enthalpies at each temperature point concentration error applied (C) Fitted direct binding enthalpies at each temperature point with heat error applied. (D) Fitted van’t Hoff binding enthalpies at each temperature with heat error applied

**Figure 5.**
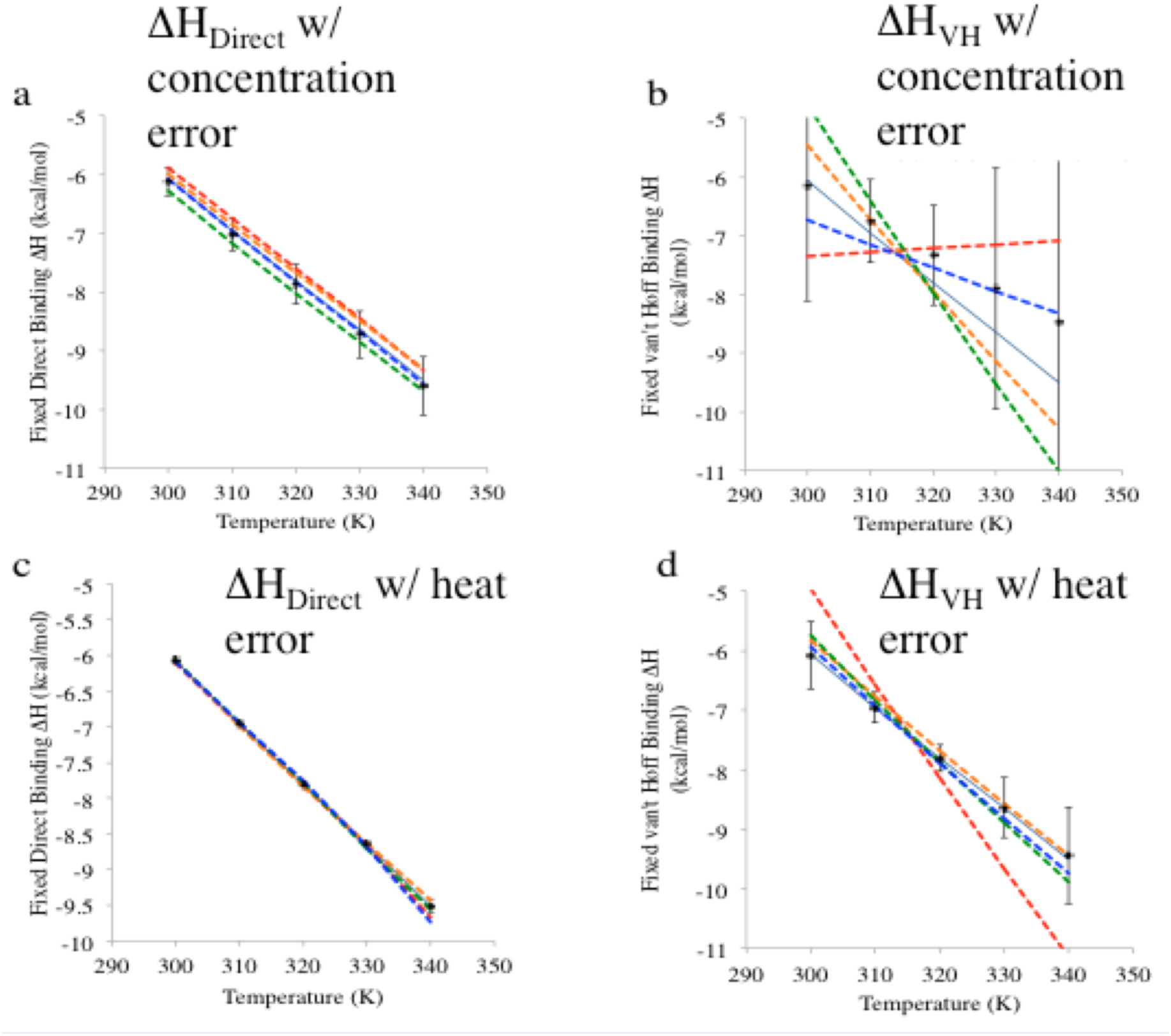
Effect of heat and concentration error on direct and van’t Hoff binding enthalpies with stoichiometry parameter N fixed at one. Dashed color lines indicate the binding enthalpies from one set of replicates; the solid line represents the nominal binding enthalpies used to generate the fitted Wiseman plots. Error bars represent one standard deviation over 1000 replicates. For concentration error, *5%* syringe and cell error was applied. For heat error, for 300-330 K points, 1% heat error and 0.13 ucal irreducible baseline error was added; for 340 K points, 3% heat error and 0.39 ucal irreducible baseline error was added. (A) Fitted direct binding enthalpy at each temperature point with concentration error applied (B) Fitting van’t Hoff binding enthalpies at each temperature point concentration error applied (C) Fitted direct binding enthalpies at each temperature point with heat error applied. (D) Fitted van’t Hoff binding enthalpies at each temperature with heat error applied.

Fixing N at its nominal value (N=1 in the present study) can often improve the reliability of thermodynamic data from ITC measurements^16,20^, so it is interesting to examine how fixing N influences error propagation into ΔH_Direct_ and ΔH_VH_. We find that, with N fixed, concentration error has a somewhat reduced effect on ΔH_Direct_ (Fig 5a vs. Fig 4a), but it now propagates dramatically to ΔH_VH_ (Fig 5b vs Fig 4b). However, heat error propagates to both values about the same with N fixed (Fig 5c,d) as with N floating (Fig 4c,d).

Finally, we examined the levels of error expected when determining van’t Hoff enthalpies under common calorimetric operating conditions, assuming the solutions are essentially ideal. This was done by using Monte Carlo sampling over heat errors and concentration errors, as previously described^16^. The resulting uncertainties in the enthalpies depend on the assumed uncertainty in concentrations, and on whether N was treated as fixed or allowed to float (Table 3), but for a typical 2% concentration uncertainty^16^, the uncertainties in van’t Hoff enthalpies run about 3-4%. This is consistent with the discrepancies D of ~2-4% in the present experimental data (above). This analysis raises the possibility that unexpectedly large discrepancies between van’t Hoff and direct enthalpies could be used to flag potentially problematic data. This idea is considered in the following subsection.

**Table 3.** Errors in calorimetric (Error_direct) and van’t Hoff (Error_VH) enthalpies in modeled data, expressed as mean unsigned error. Each row corresponds to a set of model data with the specified error applied, with the data being fit allowing N to float or N fixed, as described in the Methods. Heat errors, considered in the lower half of the table, include 1% proportional heat error at 300K and 3% heat error at 340 K, as previously determined^16^. The modeled data uses the same C values and input parameters as described in the Methods section.

|  | Syr Err | Cell Err | Error_direct<br>(%MUE) | Error_VH<br>(%MUE) |
| --- | --- | --- | --- | --- |
| **Without heat error** |  |  |  |  |
| **Float** | 2% | 2% | **1.6** | **0.1** |
|  | 4% | 4% | **3.4** | **0.1** |
|  | 6% | 6% | **4.4** | **0.2** |
|  | 8% | 8% | **6.5** | **0.2** |
| **Fixed** | 2% | 2% | **1.4** | **1.1** |
|  | 4% | 4% | **2.6** | **3.0** |
|  | 6% | 6% | **4.1** | **4.5** |
|  | 8% | 8% | **6.9** | **11.9** |
| **With heat error** |  |  |  |  |
| **Float** | 2% | 2% | **1.5** | **3.8** |
|  | 4% | 4% | **3.1** | **4.1** |
|  | 6% | 6% | **4.5** | **4.1** |
|  | 8% | 8% | **7.1** | **4.1** |
| **Fixed** | 2% | 2% | **1.2** | **3.4** |
|  | 4% | 4% | **2.9** | **4.5** |
|  | 6% | 6% | **4.4** | **6.4** |
|  | 8% | 8% | **4.8** | **10.7** |

### 3.3 Relationship between van’t Hoff consistency and magnitude of error

Given that ΔH_Direct_ and ΔH_VH_ agree perfectly with each other in the absence of experimental error, and that different sources of error result in distinct patterns of error in these two quantities, we conjectured that large inconsistencies between ΔH_Direct_ and ΔH_VH_ could be utilized to flag datasets that contain relatively high levels of error of any sort. Here we test this idea using simulated data. For these model measurements, we consistently allowed N to float in computing ΔH_VH_, because this markedly reduced sensitivity to concentration error, as noted above.

Focusing first on pure concentration error (i.e., without heat error), we find that increasing the assumed level of concentration error from 2% to 10% leads to a simultaneous increase in *E*, the deviation of ΔH_Direct_ from the true value used to generate the model data, and *D*, the discrepancy between ΔH_Direct_ and ΔH_VH_, whether N is floated or held fixed at 1 in computing ΔH_Direct_ (Fig 6). (See Eqs 5, 6 for definitions of *D* and *E*, respectively.) The same trend is observed when a realistic level of heat error is added, though only when concentration is error rises above 4% (Fig 6 c,d). Thus, for both pure concentration error and for concentration error combined with heat error, a dataset which gives good agreement between ΔH_Direct_ and ΔH_VH_ also tends to give a more accurate value of ΔH_Direct_.

**Figure 6:**
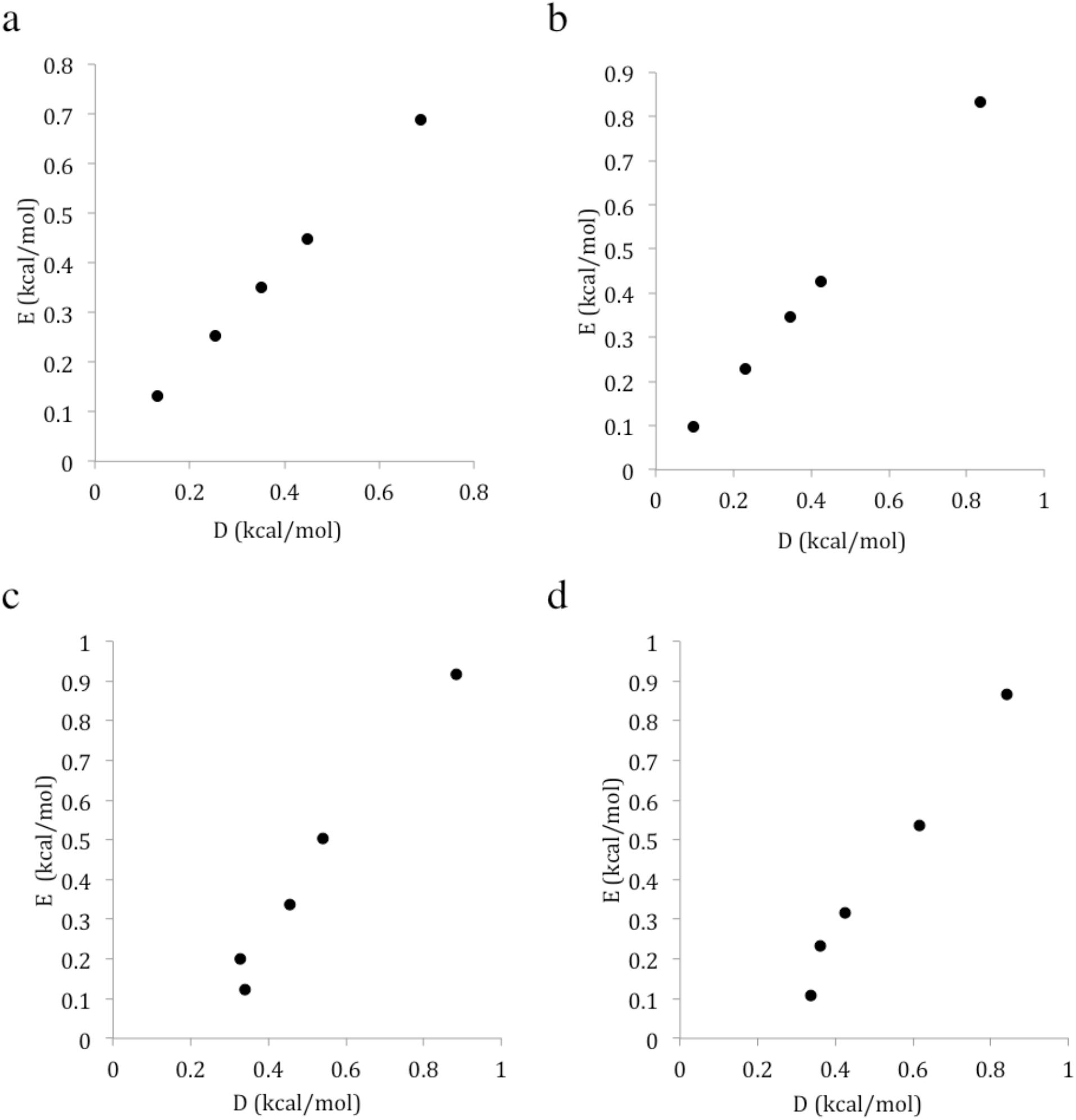
Correlation between *D*, the discrepancy between direct and van’t Hoff enthalpies and *E*, the error in fitted direct enthalpies. As points progress from bottom to top, they represent 2, 4, 6, 8, and 10% concentration error added to both syringe and cell. (A) Correlation between direct enthalpy error and consistency while N is allowed to float, with concentration error only. (B) Correlation between direct enthalpy error and consistency while N is fixed at one, with concentration error only. (C) Correlation between direct enthalpy error and consistency while N is allowed to float, with concentration error and 1% heat error added. (D) Correlation between direct enthalpy error and consistency while N is fixed at one, with concentration error and 1% heat error.

Results analogous to those in Figure 6 but with concentration error in only the syringe or cell solutions are provided in Supplementary Figures 2 and 3. These added results provide insight into how floating versus fixing N affects the relationship between the metrics *D* and *E*, through the differential effect of the treatment of N upon the propagation of cell and syringe concentration errors.^16,20^

The results above show that agreement between direct and van’t Hoff enthalpies is an indicator of data quality. Here, we extend this idea by inquiring whether this level of agreement can be used to decide whether it is better to keep N fixed or allow it to float when extracting direct binding enthalpies from calorimetric data. That is, it may be possible to discern whether floating or fixing N will decrease error in this fitted enthalpy. We used the model system used above to generate 1000 sets of five temperature points, and compared the van’t Hoff discrepancies, *D*, and calorimetric enthalpy errors, *E*, when N was floated and fixed. We then plotted *ΔE = E*_*float*_*-E*_*fixed*_ against *ΔD=D*_*float*_*-D*_*fixed*_ to look for correlation (Figure 7). Note that the van’t Hoff enthalpies here are computed based on binding free energies obtained with N floating; only the direct enthalpies were computed with either N fixed or N floated.

**Figure 7:**
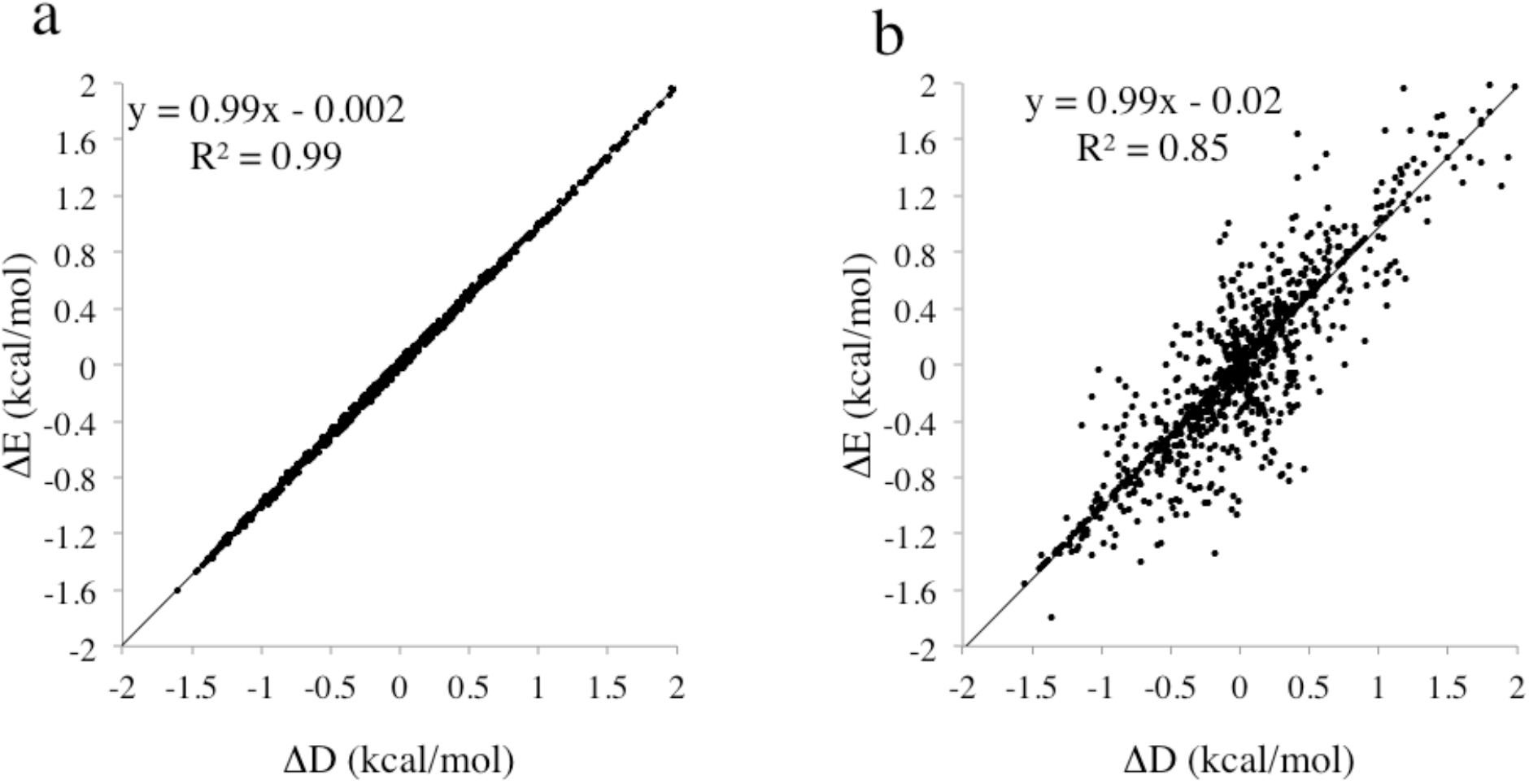
Correlation between difference in float and fix error in fitted ΔH and difference in float and fix van’t Hoff discrepancy in modeled data. Each point corresponds to a set of model ITC measurements at temperatures 300, 310, 320. 330, and 340 K, with up to 10% concentration error added to both cell and syringe and either no heat error (panel A) or with heat error based on our own experimental measurements (panel B). Here, ΔD represents the mean deviation between van’t Hoff and directly fitted enthalpies across all temperatures (Eq 5), and ΔE represents the average error in the fitted ΔH across all temperatures (Eq 6).

When zero heat error is assumed, the deviation of the direct enthalpies from the true enthalpy correlates perfectly with the discrepancy between the van’t Hoff and direct enthalpies (Figure 7a). Indeed, a linear regression of ΔE against ΔD gives a slope and correlation coefficient of essentially unity, and a y-intercept of essentially zero. In this idealized situation, then, one may determine whether to float or fix N based on which fitting method gives better agreement between the van’t Hoff and direct enthalpies (lower D). Overall, heat error (at the level estimated for the VP-ITC instrument^16^) adds scatter to the plot (Figure 7b), but the relationship between the discrepancy and error remains, with a still high R^2^. This relationship allows for a straightforward “consistency check” for measurements done over several temperatures, and may be useful in choosing whether to treat N as fixed or allow it to float, for a given dataset.

## 4. Discussion

The present study addresses a long-standing and continuing discussion in the calorimetry literature as to whether one should expect agreement between the binding enthalpy obtained directly from an ITC experiment and the binding enthalpy obtained by the van’t Hoff method. We build on prior observations of error propagation in ITC^16^,^20–22^ and expand this to include enthalpies obtained through the application of the van’t Hoff equation to binding free energies obtained with ITC. We determine the level of agreement between van’t Hoff and direct enthalpies to be expected under typical operating conditions, in the absence of significant solution non-ideality, and provide new measurements that showing agreement well within the expected uncertainties. We conjecture that the quality of agreement, compared with some prior studies, traces to two factors. The first is our use of more modern, lower-noise, instrumentation than that used in some of the older studies. The second is our use of an experimental design that avoids injecting a charged solute into the ITC cell, a practice which we suspect could generate complicating non-idealities.

Furthermore, we have shown, using model binding data, that key sources of experimental error propagate differently to the direct and the direct and van’t Hoff enthalpies, leading to discrepancies between these two fitted quantities. As a consequence, inconsistency between the direct and van’t Hoff enthalpies greater than those expected for typical operating conditions may be a useful flag for problematic data. That is, the consistency between these quantities correlates with the accuracy of the results. Similarly, one may use the level of agreement between van’t Hoff and direct enthalpies to decide whether it is preferable to treat N as fixed or to float it. The broad picture that agreement between van’t Hoff and direct enthalpy correlates with accuracy makes sense intuitively, given that, as argued below, the direct and van’t Hoff results should in principle agree to within modest uncertainties attributable to heat and concentration error. Accordingly, excessive deviations between them must be attributable to some additional error in the experiment.

The following subsections consider in more detail prior reports of large discrepancies between van’t Hoff and direct calorimetric enthalpies, consider the role of experimental error as the cause of such discrepancies, and examine prior theoretical arguments that the direct and van’t Hoff enthalpies should not in fact be expected to agree. It should be noted that the scope of this study only concerns comparison between van’t Hoff and direct enthalpies obtained from isothermal titration calorimetry. We do not consider other possible errors in van’t Hoff enthalpies obtained from other experimental methods of measuring binding affinities, such as spectrophotometry, NMR, and surface plasmon resonance.

### 4.1 Past discrepancies between direct and van’t Hoff enthalpies

In contrast with the present study, a number of prior contributions, including those of Sturtevant^5,23^, Chaires^19^, and Eggers^12^, have noted persistent discrepancies between van’t Hoff and calorimetric enthalpies and argued that the van’t Hoff equation may be flawed, or that there are other factors in play that can make this quantity impossible to obtain accurately. One reason for our different experience may be that we chose reagents which we expected would have activity coefficients near unity at the experimental concentrations, due to their low charge densities, so that our experiments would provide valid equilibrium constants. Thus, cyclodextrin is a large, neutral molecule, and the amantadine and rimantadine guests combine a bulky hydrophobic moiety with a singly charged ammonium group. That their activity coefficients were in fact close to unity is evidenced by the very low heats of dilution of the reagents (Supplementary Figure 1), and the fact that halving concentrations produced no significant change in the fitted affinity and binding enthalpy (Supplementary Figure 2). We suspect that strong nonideality was a major contributor to the van’t Hoff discrepancies seen in Eggers and Sturtevant’s studies, as they used reagents with much higher charge densities, including divalent cations and EDTA, which has four carboxylic acid groups. Binding measurements for highly nonideal solutions do not yield valid equilibrium constants, unless steps are taken to explicitly correct for the nonideality, and using invalid equilibrium constants in the van’t Hoff equation will yield incorrect estimates of the binding enthalpy. Although Sturtevant argued that nonideality should not be an issue in his experiments, this was not experimentally documented. Moreover, his measurements for a less charged pair of reagents, cyclodextrin with heptanoate, provided “puzzling” good van’t Hoff consistency^5^. Eggers’ study almost certainly was carried out under nonideal conditions, as he observed large, temperature-dependent, variations in apparent binding affinity as a function of concentration. Applying the van’t Hoff equation to these uncorrected apparent quantities could have yielded erroneous enthalpies.

Another reason for the relatively good agreement between van’t Hoff and direct enthalpies observed here is our use of an instrument that has lower heat noise than those used in some of the older studies. For example, the VP-ITC is an improvement over the Calorimetry Sciences Corporation model 4200 used by Horn et al. (JR Horn, personal communication), and indeed, they noted that the van’t Hoff discrepancies they obtained could be fully explained by instrument uncertainty^11^. Additionally, free energies measured with the VP-ITC under typical conditions have lower uncertainty than that assumed by Chaires in his modeling study of van’t Hoff disrepancies^19^. Finally, it is worth remarking that Tellinghuisen recently observed significant differences between calorimetric and van’t Hoff enthalpies, using a modern instrument, but did not argue against the validity of the van’t Hoff equation and instead speculated that the discrepancy resulted merely from problems with handling the heats of dilution^6^.

Potential explanations for apparent van’t Hoff inconsistencies are considered further in the following subsections.

### 4.2 Sources of van’t Hoff inconsistency: Experimental Error

Experimental or procedural errors are sometimes invoked to explain van’t Hoff inconsistencies. Notably, a study by Chaires found that a small curvature in the curve of binding free energy versus temperature (i.e., a small value of ΔC_p_) could be obscured by modest noise in ΔG, leading to highly inaccurate van’t Hoff enthalpies. We find that this is mostly true: noise in ΔG stemming from heat error can significantly increase the scatter of fitted van’t Hoff enthalpies, as evidenced by the results from modeled Wiseman plots. This can result from even low levels of uncertainty in ΔG, on the order of 0.05 kcal/mol. However, given the modest levels of heat error in the VP-ITC instrument, obtaining van’t Hoff enthalpies in good agreement with direct enthalpies should not be as difficult as previously predicted, even with smaller ΔC_p_’s. The value of ΔC_p_ used for the model data in sections 3.2 and 3.3 was −85 cal/mol/K, and the ΔC_p_ for both experimental datasets (which gave reasonable consistency) were ~-70 cal/mol/K, suggesting that, even if ΔC_p_ is relatively small (Chaires’ study assumed ΔC_p_ of 200 cal/mol/K), good consistency should still be achievable, at least in a well-controlled experimental setting.

We also find that concentration error has minimal effect on van’t Hoff enthalpies, in contrast to the known sensitivity of the direct calorimetric enthalpy to concentration error^16,20,22,24^. This is presumably because we use the same solutions across for the full temperature series of experiments, so that any concentration error is constant across the series. Thus, any concentration error that is present shifts the apparent binding affinity at each temperature point by about the same amount, allowing the slope and curvature of the free energy versus temperature curve to remain essentially unaffected, and hence preserving the van’t Hoff enthalpies. In contrast, if different solutions were used at the different temperatures, we expect that van’t Hoff enthalpies would be greatly affected by concentration error in much the same way that they are affected by heat error, where the free energy at each temperature has independent scatter that affects the curvature and thus the binding free energies. The net effect would be a marked increase in the discrepancy between direct and van’t Hoff enthalpies. It is therefore essential that measurements of van’t Hoff enthalpies utilize the exact same solutions for the measurements at all temperatures. We guess that previous studies comparing direct and van’t Hoff enthalpies used the same solutions across all measurements, but this methodological detail is rarely specified, so the use of different solutions at the different temperatures might, in some cases be an added source of discrepancy.

As noted previously^6,21^, the van’t Hoff equation applies to standard thermodynamic quantities, which pertain to ideal solutions, so apparent deviations between direct and van’t Hoff binding enthalpies can arise if the solutions used deviate significantly from ideality, unless pains are taken to correct back to standard state. Carrying out such corrections would appear to require knowledge of the activity coefficients of the reactants and product, both alone and as mixtures, and perhaps also knowledge of the dependency of the partial molar enthalpies of these species upon concentration. These quantities are often not available, so corrections cannot easily be made. However, as previously emphasized^21^, one may at least determine whether the binding free energy and enthalpy are insensitive to concentration at the desired experimental conditions, as done in the present study.

In the present study, then, the greatest contribution to inconsistency between van’t Hoff and direct enthalpies seems to be the heat error intrinsic to isothermal titration calorimetry instruments. Despite the inevitable presence of these errors, given a robust C value across the temperature range (such as the range of 5 to 50 used in the present models), and based on current estimates of heat error^16^, we anticipate that consistency should be achievable to within about 4-5% mean unsigned error between van’t Hoff and calorimetric enthalpies. Given that earlier ITC instruments had larger heat errors than the instrument used here, some past reports of van’t Hoff inconsistency likely stemmed from this class of experimental error.

### 4.3 Sources of van’t Hoff inconsistency: Theoretical

Theoretical explanations have also been proposed to explain apparent inconsistencies between direct and van’t Hoff binding enthalpies. Thus, Weber argued that the van’t Hoff equation is fundamentally wrong, due to confusion between the entropy change of the reactants themselves and the entropy change of the whole system^7,8^, but this line of reasoning was convincingly rebutted^9,10^ and is no longer current in the literature. Another theoretical argument came from Sturtevant, who suggested that solvent effects play a role in discrepancies between van’t Hoff and calorimetric enthalpies^4,5^. In this view, the rearrangement of solvent around the solutes on binding leads to an additional free energy term not accounted for when calculating dΔG°/dT to generate van’t Hoff enthalpies. Castellano and Eggers^12^ have recently elaborated this idea, arguing that the standard binding free energy comprises not only the customary difference in standard chemical potentials of the reacting solutes (e.g. a protein, a ligand, and their complex), but also an added contribution, termed the desolvation free energy, from the change in chemical potential of solvent molecules released from the binding surfaces to bulk upon binding. They suggest that this term is missing from conventional treatments of the thermodynamics of reactions in solution, and that this concept can be used to explain many experimental observations of discrepant direct and van’t Hoff enthalpies.

We disagree with this view, and now present a brief statistical thermodynamic derivation of the van’t Hoff equation, which shows that this relation holds true even when solvent is fully accounted for, and that changes in solvation of the reactants on binding do not require any modification to solution binding theory. We note that, although the van’t Hoff equation can be derived based purely on thermodynamics the statistical thermodynamics perspective is informative here because it makes the roles of the various molecular species explicit.

The standard free energy of reaction is the difference between the standard chemical potentials of the product and reactant species^25^, which here are the bound and free species^26^. This is because the standard chemical potential of a molecular species is the change in free energy of the system when one mole of the species is added to the system. When the reaction proceeds forward by one mole under standard conditions, one mole of each reactant is consumed and one mole of product is generated. Consequently, the change in the free energy of the system is the difference in chemical potentials:

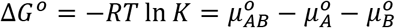

Here 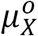 is the standard chemical potential of species X = AB, A or B, where A and B bind to form the complex AB. Note that the standard chemical potentials pertain to hypothetical ideal solutions at standard concentration (typically 1M), and that *K* is a concentration-independent quantity, by definition. Because no water is added or consumed in the course of the reaction, the chemical potential of water does not enter explicitly. Instead, the presence of water enters through its influence on the chemical potentials of A, B, and AB. Thus, the standard chemical potential of species X is given by^14,26^

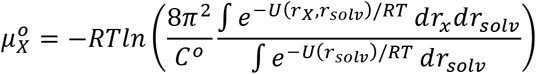

Here *8π*^2^ accounts for the free rotation of the solute, C° is the standard concentration (typically 1 M), r_x_ represents the internal coordinates of species X, r_solv_ represents the coordinates of the water and other solvent molecules in the coordinate frame of the solute, U(r_X_,r_solv_) represents the potential energy of the solute-solvent system as a function of these coordinates and U(r_solv_) is the potential energy of the solvent in the absence of the solute. (See refs 17 and 18 for details.) This expression accounts for all favorable and unfavorable interactions of water with the solute, so the difference in chemical potentials in Δ*G°* fully accounts for the fact that the bound complex has less solvent-exposed surface than the unbound reactants. Now, recognizing that

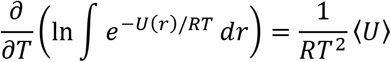

and using the equations above, it is straightforward to complete a derivation of the van’t Hoff equation^14^:

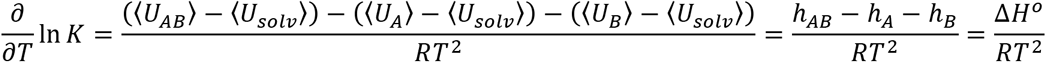

Here, we have used the fact that h_x_, the standard partial molar enthalpy of solute X, is the change in mean energy when a mole of solute is added to pure solvent, with the simplifying approximation that any net volume change is negligible. Thus, the van’t Hoff relation follows directly from the statistical thermodynamics of ideal solutions, and it fully accounts for the changes in solute-water interactions that occur on binding.

We therefore see no basis for the view of Castellano and Eggers that changes in reactant solvation on binding can lead to violations of the van’t Hoff equation. Furthermore, the chemical potential of water, which is defined as the change in the free energy of the system when another mole of water is added, is not a function of position. Therefore, we do not believe there is a sound theoretical basis for their claim that water at the solute interface has a different chemical potential from bulk water in the same flask. Analogous reasoning holds for other components of the system, such as protons. Thus, Horn et al. showed that linked equilibria should not affect van’t Hoff enthalpies, and therefore can be discounted as a possible explanation for discrepancies between van’t Hoff and direct enthalpies^27^.

If the reworking of solution thermodynamics proposed by Castellano and Eggers does not hold, then another explanation must be sought for the apparently severe discrepancy between van’t Hoff and direct enthalpies in their EDTA-Ca ITC experiments. They observe good agreement between the direct and van’t Hoff enthalpies in experiments where the reactants are at low concentration, but increasing inconsistency with rising concentrations. This pattern suggests a role for non-ideality, which also tends to become more marked with increasing concentration. Non-ideality can play a role because the equilibrium constant which enters the van’t Hoff equation is, by definition, independent of concentration, whereas Castellano and Eggers may instead have used an apparent equilibrium constant, by which we mean the equilibrium ratio of concentrations. When solutions are not ideal, this is a concentration-dependent quantity, and there is no reason to expect the van’t Hoff equation to work if it is used in place of the equilibrium constant itself. In the present instance, we anticipate that the highly charged reactants, Ca^2+^ and EDTA^2−^ or EDTA^3−^, will have substantially lower activity coefficients at high concentration than at low concentration. However, the calcium-EDTA complex will likely have an activity coefficient near unity, because its net charge is near zero. Thus, raising the concentrations of all species will lead to an apparent weakening of binding, as was observed experimentally. For example, if at high concentration the activity coefficients of the reactants are *γ*_*C*_= *γ*_*E*_= 0.5, while that of the complex is *γ*_*C.E*_= 0.9, then, given the usual expression for the equilibrium constant,

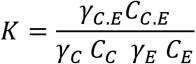

the apparent equilibrium constant will be

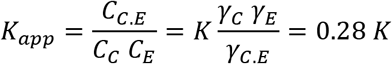

Thus, the reduced activity of the reactants vs. the products, at high concentration, leads to less binding. Although one may insert K_app_ into the van’t Hoff equation in place of K and thus obtain an apparent van’t Hoff enthalpy that differs from the direct binding enthalpy, this inconsistency is not in fact an example of van’t Hoff inconsistency, because the van’t Hoff equation by definition uses K, not K_app_. Additionally, the partial molar enthalpy of a solute may change with concentration, which would be another source of discrepancy in observed calorimetric and van’t Hoff enthalpies. It is also worth emphasizing that the large change in the apparent equilibrium constant with concentration observed in the EDTA-Ca experiments is itself direct evidence that the solutions are not behaving ideally at high concentration. Thus, nonideal solution behavior likely explains the large apparent van’t Hoff inconsistency seen by Castellano and Eggers for their measurements at high concentration, and their results do not imply that the long-standing theory of solution thermodynamics needs to be revised. Nonideality might also help explain other cases of apparent van’t Hoff inconsistency, as previously emphasized by Pethica^21^, particularly in cases where one or more of the reactants is highly charged.

Given the apparent occurrence of significant nonideality in prior calorimetric studies of small molecules, it is of interest to consider whether nonideality also comes into play in biomolecular applications. We are not aware of many measurements of protein activity coefficients, but a study of the protein alpha-chymotrypsin reported activity coefficients within a few percent of 1 for concentrations up to about 1 *µ*M, under various salt conditions^28^. Thus, applications of ITC that use protein concentrations in this range or lower probably avoid serious nonideality, at least for the protein solute. The concern is greater, however, for highly charged solutes, such as nucleic acids. The presence of nonideality may be checked by looking for concentration dependence of the results, or by comparing van’t Hoff and direct enthalpies, because, as mentioned above, the concentration dependent apparent K and partial molar enthalpy would cause discrepancies between the two enthalpies.

### 4.4 Using van’t Hoff consistency to check for error

Given that the direct and van’t Hoff enthalpies should, in principle, be in accord, an apparent discrepancy between these two quantities beyond that expected based on known experimental uncertainties can be a useful flag for the presence of experimental errors of various types, such as errors in solution concentrations, unexpectedly high heat errors, or marked nonideality of the solutes. This concept may be particularly useful in settings where high accuracy is desired. In addition, the van’t Hoff consistency check can be used to guide the decision of whether to fix or float the stoichiometry parameter, N, in fitting ITC data, as our model calculations show that greater consistency correlates with greater accuracy.

## 5. Conclusions

This study yields several key observations regarding the nature of enthalpies obtained from both calorimetry (direct) and van’t Hoff methods using an isothermal titration calorimeter:

- The direct and van’t Hoff enthalpies should in principle agree well
- Inconsistency between the van’t Hoff and direct enthalpies can results from concentration error, heat error, or solution nonideality.
- Inconsistency between the van’t Hoff and direct enthalpies is a sign of experimental error.
- Concentration error has little effect on van’t Hoff enthalpies when N is floated, given that the same solution is used for experiments at all temperatures.
- Concentration error can propagate strongly to direct enthalpies, with cell error propagating when N is fixed, and syringe error propagating when N is floated.
- Heat error propagates strongly to van’t Hoff enthalpies, whether or not N is allowed to float.
- Heat error does not propagate strongly to direct enthalpies, at least for the C values considered here.
- The decision to float or fix N can be guided by examining the discrepancy between direct and van’t Hoff enthalpies with both N float and N fixed: a lower discrepancy correlates to a lower error relative to the true enthalpy.

## Acknowledgements and Disclosures

M.K.G. has an equity interest in, and is a cofounder and scientific advisor of VeraChem LLC. S.A.K. acknowledges training support from the Molecular Biophysics training grant (T32 GM008326). This publication was supported by the National Institute of General Medical Sciences of the National Institutes of Health (NIH) Grant GM61300. These findings are solely of the authors and do not necessarily represent the views of the NIH.

